# Robustness in Hopfield neural networks with biased memory patterns

**DOI:** 10.1101/2024.10.31.621250

**Authors:** Ali Montazeri, Robert Schmidt

## Abstract

Biological neural networks are able to store and retrieve patterns in the presence of different types of noise. Hopfield neural networks have been inspired by biological neural networks and provide a model for auto-associative memory patterns. An important parameter in these networks is the pattern bias, i.e. the mean activity level of the network, which is closely related to the degree of sparseness in the coding scheme. Here we studied the relation between robustness against different types of biologically-motivated noise and pattern bias. To do so, we developed performance and robustness measures, which are applicable to varying degrees of sparseness of the memory representations, using analytically-optimized thresholds and corruption tolerances adjusted by mutual information. We then applied these tools in numerical simulations and found that, for different types of noise, the pattern load, i.e. the number of patterns that the network has to store, determined which pattern bias is most robust. Across different types of noise, the higher the pattern load was, the more biased was the most robust performing pattern representation. Given the variation in the sparseness level in different brain regions, our findings suggest that memory pattern encoding schemes (i.e. degree of sparseness) in different brain regions might be adapted to the expected memory load in order to best mitigate the adverse effects of disruptions.

## 1 Introduction

One characteristic of memory in biological organisms is the robust performance under noisy conditions (Rolls & Deco, 2010). Hopfield neural networks have been inspired by biological neural networks and provide a model for auto-associative memories (Hopfield, 1982). The effect of noise on retrieval and capacity in Hopfield networks has been of high interest already in the early seminal studies of the Hopfield network properties (Amit et al., 1987), and is relevant as a computational model of memory systems in biological organisms. Furthermore, understanding robustness against noise is relevant for neurological disorders such as Alzheimer’s disease, in which, during the preclinical phase, memories seem to endure pathological alterations (Long & Holtzman, 2019).

Noise can be introduced in Hopfield networks on different levels. In analogy to statistical mechanics of physical systems, a common model for noise is to consider thermal fluctuations of the activation function, whose strength is described by a temperature parameter. As a model for auto-associative memories in the brain, the temperature parameter, or an equivalent stochastic threshold (Hertz et al., 1991), captures some aspects of the random variability in neural activity. However, in biological neural networks there are also other types and sources of noise (Faisal et al., 2008). For example, structural noise includes random variation of the connection weights (e.g. due to variations in post-synaptic potentials; Ribrault et al., 2011), but also random loss of connections or whole neurons (e.g. due to injury or neurodegenerative diseases). In addition, the initial pattern presented to the network for retrieval can be subject to noise, so that for a random subset of units the states are flipped when a stored pattern is presented. Optogenetic methods in neuroscience (Boyden et al., 2005) also motivate to consider a variation of this type of noise, so that a random set of units is just collectively activated or deactivated, irrespective of the actual pattern. From a biological perspective, Hopfield networks, as a model for auto-associative memories in the brain, should exhibit robustness against the biologically-motivated types of noise.

In addition to noise, also the coding schemes employed by Hopfield networks have been studied especially in the context of memory capacity. In the classic Hopfield network memory patterns are unbiased, i.e. that a given pattern activates 50% of the network nodes. Subsequent work demonstrated the ability of Hopfield networks to also store biased patterns, i.e. patterns that activate less than 50% of the network nodes (Amit et al., 1987). Biased, or sparse, patterns are also common in biological neural networks, and it turns out that also Hopfield networks can increase their storage capacity for biased patterns (Tsodyks & Feigel’man, 1988). However, dealing with biased patterns requires a modification of the Hopfield network learning rule as biased patterns otherwise increase spurious states and thereby degrade performance. Another aspect of network performance that goes beyond storage capacity is robustness.

The robustness of a Hopfield network relates to the dynamical properties of the network. In the energy landscape the desired stable states form basins of attraction. The relative size and energy levels of the basins of attraction of the desired stable states in comparison to those of the spurious attractors determine the chances for a successful memory retrieval (Rolls & Deco, 2010). This is schematically illustrated in Figure 1A&B, where the storage capacities are similar but the attractors associated with the memory states in A have larger and deeper basins of attraction than in B. Therefore, even when starting from the desired stable states, the behaviour of the two networks will be different in the presence of noise.

**Figure 1:**
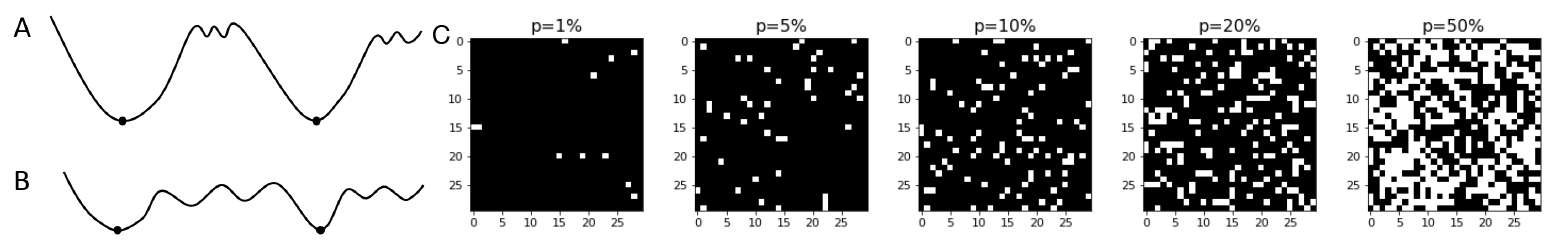
Energy landscapes and pattern bias. (A, B) Schematic illustration of the energy landscape of two networks with differences in the size and depth of the basins of attraction. The memory states’ attractors are shown by dots. When starting in a stable state, noise might push the network in (B) into a local minimum, while network in (A) would remain in the desired state. (C) Examples of memory pattern configurations from very sparse (left) to unbiased (right) representations. The network has 900 (30×30) nodes. The white and black squares represent 1 and 0 states, respectively.

Here we propose and examine a measure of robustness in Hopfield networks to store biased and unbiased patterns. We found for different types of noise that the most robust pattern bias depended on the pattern load. This indicates that there is a trade-off between memory load and robustness so that the memory system should employ a coding scheme, i.e. a level of pattern bias, that depends on the expected pattern load. We propose that this might be relevant for biological networks, so that coding schemes employed in different brain regions might be tuned towards optimal performance based on the expected memory load in those regions.

## 2 Methods

### 2.1 The Hopfield network

The attractor neural network proposed by Hopfield (Hopfield, 1982) accounts for several characteristics of auto-associative memories including pattern completion and memory retrieval from an incomplete or flawed clue. In this network, the *N* nodes are McCulloch-Pitts neurons, which are either in a (quiescent) 0 state or in an (active) 1 state. For each neuron *i*, given the symmetric connection strengths *W*_*ij*_ from neurons *j* to *i*, the state variable *V*_*i*_ is updated by the following dynamical rule

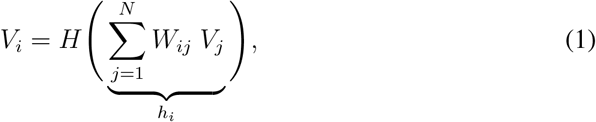

where *h*_*i*_ is the neuron’s local field, and *H* is the Heaviside step function with the value of 0 for negative arguments and 1 for positive arguments. In the classic model the update is carried out asynchronously, so that units are updated one at a time. The goal is to store a set of *M* binary input patterns *{ξ*^*µ*^, 1 ≤ *µ* ≤ *M }* via adjustments to the connection weights *{W*_*ij*_, 1 ≤ *i, j* ≤ *N }*, such that the desired memories become the network’s stable states. The weights are determined based on Hebbian correlation learning, i.e.

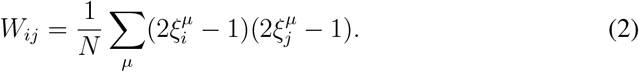

In standard Hopfield networks there are symmetric recurrent connections between all nodes (but no self-coupling, i.e. *W*_*ii*_ = 0).

For unbiased patterns, each 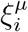 is a (fixed) random and independent Bernoulli bit with a mean level of activity *p* of 50%, i.e.

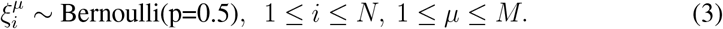

The randomness and independence of bits bring mathematical convenience for studying the network’s characteristics, for instance in the signal-to-noise analysis of the model.

An important contribution of Hopfield (1982) was to introduce the notion of an energy function into neural networks:

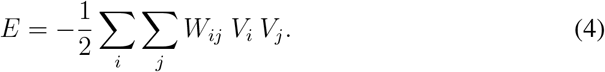

The activity update dynamics of the system ensure that the energy function always decreases or remains constant. The stable states or attractors of the network are the local minima of the energy surface, and the “basins of attraction” correspond to the valleys around each minimum (Hertz et al. 1991; Figure 1A&B).

For the stored patterns to be retrieved successfully from an incomplete or distorted stimulus, they must coincide with the network’s attractors (i.e. the local minima of the energy function). Thereby, an initial network configuration, with bit states being different from the original pattern, is updated step by step towards the stored pattern, where no further changes take place. However, the energy landscape has also local minima corresponding to “spurious” states, that can interfere with the memory recall.

The storage capacity of the network is the maximal number of stored patterns *M*, for which a retrieval is possible. The storage capacity of a Hopfield network can be expressed using the load parameter *α* = *M/N* (MacKay, 2003) that takes into account the network size *N*. Small deviations between the stored and retrieved pattern are considered acceptable and increase the storage capacity considerably (MacKay, 2003). To measure the distance between a stored and a retrieved pattern we use the (normalized) Hamming distance, which calculates the ratio of bits that are different between two binary configurations ([0, 1]).

### 2.2 The Hopfield model for biased patterns

The original Hopfield model works well for uncorrelated patterns when the Bernoulli probability *p* in Eq. 3 is 50%. Memory patterns are called biased if the mean neural activity *p* in Eq. 3 deviates from 0.5, and we refer to them here as sparse if *p* ≪ 0.5. Schematic examples of memory patterns with different levels of bias are shown in Figure 1C.

In order to store biased patterns, the Hopfield model must be modified to avoid spurious states from becoming dominant. The learning rule is adjusted as follows (Amit et al., 1987):

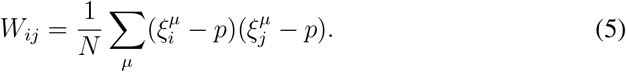

In addition, the Heaviside step function in the dynamical rule (Eq. 1) is replaced by an optimized activation threshold (the same for all nodes), which keeps the capacity in the order of 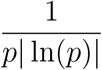 as *p* → 0. (Gardner, 1988; Tsodyks & Feigel’man, 1988; Vicente & Amit, 1989).

For a given *p*, the optimized activation threshold can be obtained by a signal-to-noise analysis of the network (Vicente & Amit, 1989). Given that the network state is a desired memory pattern *υ*, i.e.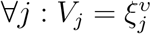, the local field *h*_*i*_ can be decomposed into a signal part, which reinforces the desired memory, and a remaining quasi-random cross-talk noise due to interference from other stored patterns. The derivation of the signal and noise parts is provided in Appendix A (see also Buhmann et al., 1988). Assuming a uniform activation threshold *θ*, the derivation results in the local field *h*_*i*_ to have the signal parts:

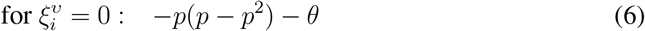

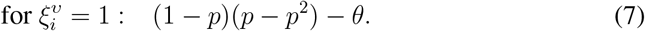

The cross-talk noise has a zero mean with the variance:

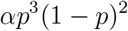

Hence, the optimal threshold corresponds to the average of the expressions (6) and (7) being zero, which yields:

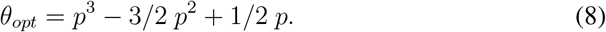

The signal and noise parts are shown schematically in the Figure 2A as mean values and standard deviations of the normally-distributed local fields, respectively. The optimal threshold shifts the two distributions, so that the absolute value of their means is identical (see example in Figure 2B). The optimal threshold has a concave shape as a function of *p* and there are upper and lower bounds for *θ* to keep the averages for 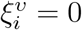 and 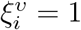 on their current side of the x-axis (Figure 2C).

**Figure 2:**
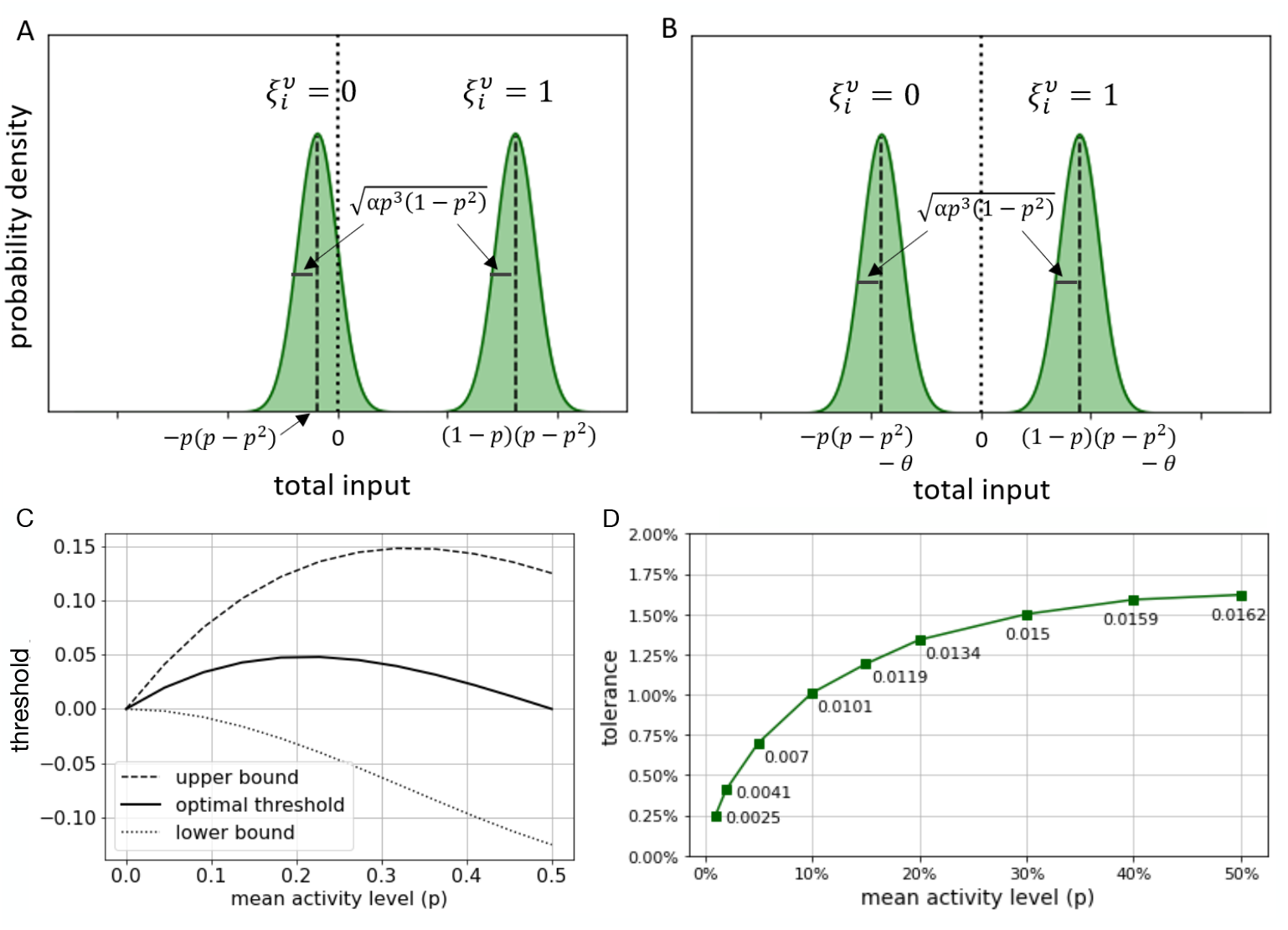
Optimal thresholds and tolerance for different pattern bias. (A, B) The probability density functions for the local field *h*_*i*_ are given by the signal part (the mean of the Gaussians; dashed black lines) and a cross-talk noise part (the standard deviation), when the bit 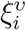 of the stored pattern is either 0 or 1. Without an activation threshold (A) the mean lies below 0 for 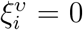, but due to the noise part the distribution extends to positive values, which would lead to an erroneous activation according to Eq. 1. Introducing an optimal activation threshold *θ* shifts the means of the distributions so that the unit is active only for 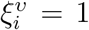 and not for 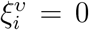 (B). (C) The optimal threshold varies for different levels of mean activity. The upper and lower bounds of the thresholds ensure that the means of the distributions (B) do not have the same sign. (D) For different levels of pattern bias, tolerances, i.e. corruption levels (in Hamming distance) that corresponded to 88% normalized mutual information, were determined.

The capacity of the original Hopfield network increases by a factor of 2 if the so-called S-variables ∈ *{±*1*}* instead of V-variables ∈ *{*0, 1*}* are used (Hopfield, 1982). Given the linear transformation between the two representations, i.e. *S*_*i*_ = 2*V*_*i*_ − 1, the S-model is equivalent to the V-model with the random thresholds 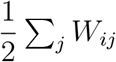 Similarly, for the biased memory patterns, if the “random local fields” *p* Σ _*j*_ *W*_*ij*_ are added to each node *i* of the V-model, the variance of the cross-talk noise improves from *αp*^3^(1− *p*)^2^ to *αp*^3^(1 − *p*)^3^ (see appendix B for details). The application of this random local threshold is equivalent to the optimized model from a continuous family of models discussed by Vicente and Amit (1989). In summary, the resulting optimal signal-to-noise ratio with both a uniform external field *θ*_*opt*_ which optimizes the signal, and the random local fields *p* Σ_*j*_ *W*_*ij*_ which correlate with the patterns and reduce the cross-talk noise is

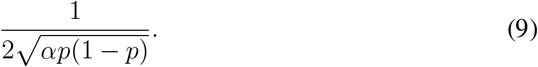

Thus, as expected, the recalling behavior improves for sparse patterns. The modified dynamics of our model are

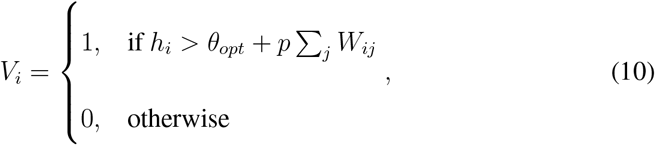

with the following energy function (Vicente & Amit, 1989)

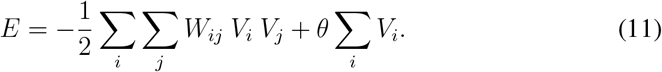

### 2.3 Noise in the network

The conventional way to add noise to the neural network (or temperature to the Ising model) is to make the update rule for units stochastic by introducing the logistic function *f*_*β*_, so that an activation state *V* is taken with probability *P* :

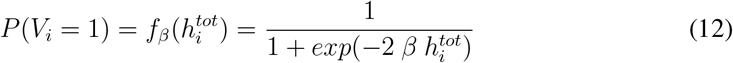

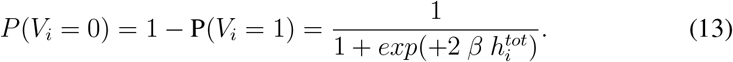

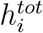 is the total field equal to *h*_*i*_ − *θ* − *p*Σ _*j≠i*_ *W*_*ij*_ for the dynamics in Eq. 10, and the parameter *β* relates to the pseudo-temperature *T* by

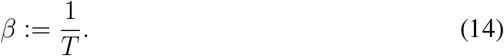

The parameter *T* controls the steepness of the sigmoid function near 0. The above-mentioned noise is equivalent to adding a stochastic threshold *θ*_*stoch*_, drawn from the probability density 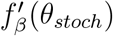, to the deterministic model (Hertz et al., 1991).

The ability of Hopfield networks to robustly retrieve stored patterns and work as a memory device depends on the regime in the T-*α* plane. For a given temperature, networks with small load parameters have attractors associated with the stored memory patterns. In this regime spurious states still exist, and can take the form of mixture states (Amit et al., 1985a) or states uncorrelated to to any of the stored patterns. If the load parameter or temperature is increased, those states start to dominate the network. While there is a temperature range in which the efficiency of the system is improved (Amit et al., 1985a), the network becomes no longer useful as a memory device if, given a temperature *T*, a critical load parameter *α*_*c*_(*T*) is exceeded, or if the temperature is increased to an overly high value (Hertz et al., 1991). The critical load parameter in the noiseless original Hopfield network *α*_*c*_(0) was found to be almost 0.138 (Amit et al., 1985b), and increases superlinearly for a model with biased patterns and the dynamics of Eq. (10) (Vicente & Amit, 1989).

While the noise in the form of the logistic function and temperature as in Eq. (12) is relevant in statistical mechanics, there are other types and sources of noise in biological neural networks (Faisal et al., 2008). Therefore, we applied the following different types of noise separately to the network to investigate the model’s robustness against them:

A. Stochastic threshold. A stochastic threshold was added to the model as described above. The pseudo-temperature in the logistic function of Eq. 12 ranged from 0 to 1*/*8, which corresponds to a range of 0 to 1 using S-variables.
B. Weight noise. Weight noise was implemented by adding Gaussian noise to the weight matrix. This additive noise was generated from the normal distribution with a standard deviation between 0 (no weight noise) and 0.002. For comparison, the highest standard deviation that was used corresponds almost to the standard deviation in the weight matrix of the model with unbiased patterns, which was the highest among the different levels of *p*. The self-coupling terms *W*_*ii*_ were still set to 0.
C. Connection dilution. Connection dilution was incorporated in the model by setting a random set of weights in the weight matrix to 0. The probability of diluting an individual, recurrent connection ranged from 0% to 100%.
D. Node dilution. For node dilution a random set of nodes was completely removed from the model. The fraction of deleted nodes ranged from 0% to 90%. Thereby, in few cases a pattern may even be completely removed, or no longer distinguishable from another one, so its retrieval failed automatically.
E. States turning active. To implement this new type of noise a random set of states of a stored memory was set to 1, irrespective of whether they were 0 or 1 in the stored pattern, to produce the initial configuration for the simulation. A range of 0 to *N* states are chosen for this distortion.
F. States turning quiescent. In correspondence to E) this noise type was implemented by setting a random set of states of a stored memory 0 to produce the initial configuration for the simulation. A range of 0 to *N* states were chosen for this distortion.
G. State flipping noise. Finally, this commonly used type of noise was implemented by flipping a random set of states of a stored memory from 0 to 1 and vice versa to produce the initial configuration for the simulation. A range of 0 to *N* states are chosen for this distortion. Since *n* flips of the states of a memory pattern results exactly in *n/N* Hamming distance, this form of distortion is widely used in the literature.

### 2.4 Patterns

The number of nodes *N*, or pattern size, was set to 5000. A set of *M* random patterns with independent Bernoulli bits was generated, each with a mean level of activity *p* determining the pattern bias:

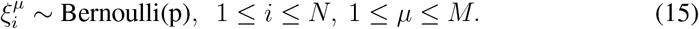

The pattern bias, or sparseness level, given by *p* ranged as follows (with 50% corresponding to unbiased patterns and 1% to extremely sparse patterns):

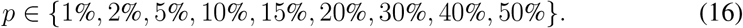

The number of patterns to store was also systematically varied with the load parameter *α* ranging from 5% to 50%.

### 2.5 Simulation procedure

For each choice of *p* and *α* and for each experiment, *M* random patterns were generated, and the weight matrix was determined according to the modified Hebbian learning rule (Eq. 5). The optimal uniform threshold *θ*_*opt*_ was set according to Eq. 8 and the random local fields *p* Σ _*j*_ *W*_*ij*_ were then obtained accordingly. Next, each type of noise was investigated separately. For noise that affected the network dynamics (types A-D, see above), the network was initialized with one of the memory patterns at a time. For the other noise types (E-G) the network was instead initialized using the distorted versions of the memory patterns. In all cases synchronous dynamics were used, so that all units were updated simultaneously, as there is no significant change of results in comparison to the asynchronous updating (Hertz et al., 1991), but an enormous computational advantage. The maximum number of update steps was set to 50, which was sufficient to reach a stable state. Afterwards, the Hamming distance between the original pattern and the settled network’s state was calculated. For the case of node dilution, the Hamming distance was calculated based on the remaining nodes. For each type and magnitude of noise, 100 experiments were performed. To keep the runtime of the relatively large networks feasible, PyTorch Tensor computing with a graphics processing unit (GPU) acceleration was used.

### 2.6 Performance and Robustness measures

The performance of Hopfield networks can be assessed based on the critical value *α*_*c*_(*T*), for which there is a discontinuous jump in Hamming distance between stored and retrieved patterns. For a network just below this critical value and with unbiased patterns, the Hamming distance between a desired memory and its associated attractor is 1.6% (MacKay, 2003). This value is called “corruption tolerance” for a successful memory retrieval, and depends on the pattern bias. To obtain a comparable tolerance level for the different values of the pattern bias, the normalized mutual information between memory patterns and their corrupted versions was used. This is necessary for the comparison because the total information is maximized when *p* = 50% and reduces as the pattern representation becomes sparser. The normalized mutual information between patterns *X* and *Y* determines the fraction of information in X that is contained in Y (Cover & Thomas, 1991), and is given by:

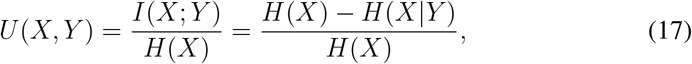

where *H*(*X*) is the entropy and *H*(*X*|*Y*) the conditional entropy. In our case *X* and *Y* can be considered as two random variables, which assume the value 0 or 1. Thus, there are four possible events *X* = *i, Y* = *j* with probabilities *p*_*ij*_ that sum to 1. The two marginal distributions are given for *X* by *p*_*X*=0_ = *p*_00_ + *p*_01_ and *p*_*X*=1_ = *p*_10_ + *p*_11_, and for *Y* by *p*_*Y* =0_ = *p*_00_ + *p*_10_ and *p*_*Y* =1_ = *p*_01_ + *p*_11_. Hence, the general definitions of *H*(*X*) and *H*(*X*|*Y*) reduce to

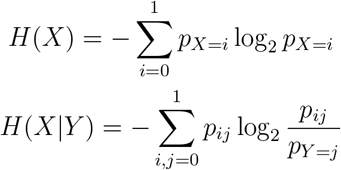

To find the relationship between the level of corruption (in Hamming distance) and the normalized mutual information, numerical simulations were used. For unbiased patterns, the mutual information associated with around 1.6% error was 0.88 bits. Therefore, we used this value for calibration purposes to find tolerances yielding comparable corruption levels for biased representations (Figure 2D). These tolerances were then used to measure the network’s performance as the “hit ratio”, i.e. the ratio of retrieved patterns with a smaller Hamming distance than the tolerance level.

The robustness measure was based on the relation between noise levels and hit ratio. In general, increasing noise levels eventually lead to a break down of the network performance and the hit ratio rapidly drops to zero. For each type of noise, the robustness was therefore measured by calculating the area under the curve (AUC) of the hit ratio versus the entire noise range, normalized by the maximum possible area. The normalized AUC is a non-parametric measure which aggregates the performance of the network against a certain noise type.

## 3 Results

The goal of our study was to determine the relation between the robustness of the Hopfield network and pattern bias for varying memory loads. To do so we first confirmed the optimality of the analytically-derived thresholds, which then were used in the subsequent simulations. Next, we examined the networks’ load capacity in terms of the hit ratio and the Hamming distance, which form the basis for our robustness measure. Finally, we applied our robustness measure to test how different types of noise affect the network performance depending on the pattern bias.

### 3.1 Optimal activation thresholds

To calculate the optimal activation thresholds we used the signal-to-noise ratio analysis (Eq. 8; see also Figure 2). We verified this approach numerically by running simulations for different values of thresholds *θ*, for each level of pattern bias. We found that the analytically-derived optimal thresholds indeed coincided with the maximum hit ratio of 1 in all cases, as exemplified in Figure 3A for *α* = 0.1 (i.e. 500 patterns). The range of thresholds for which the network showed highest performance decreased for sparse patterns, as expected from the decreasing distance between the upper and lower bounds (Figure 2C).

**Figure 3:**
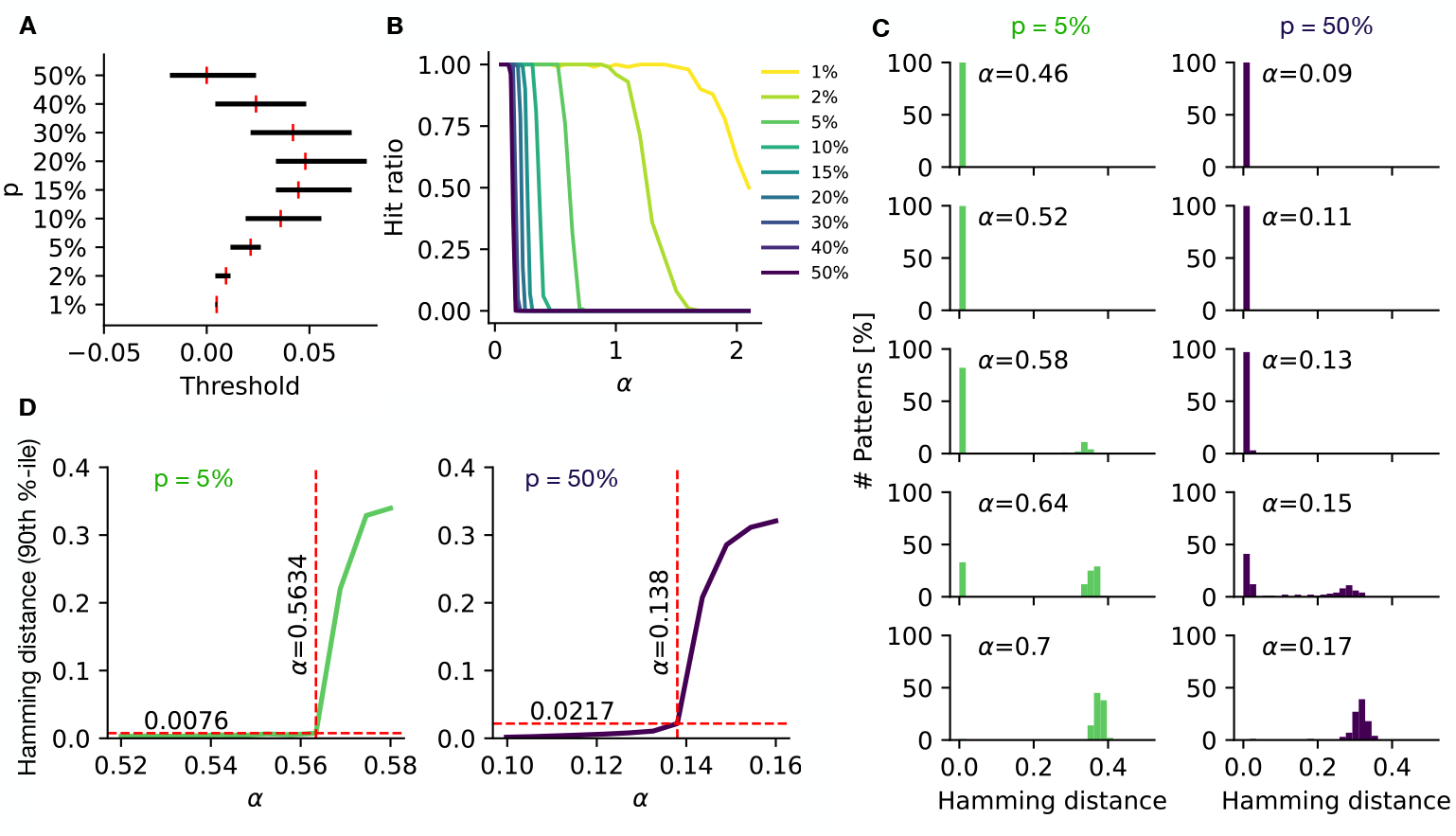
Network performance and capacity. (A) For different levels of pattern bias (y-axis), the performance of different thresholds was tested in numerical simulations (here *α* = 0.1). The black lines indicate the range of thresholds for which the hit ratio was 1 (outside this range the hit ratio quickly degraded to 0). The red vertical lines indicate the analytically-derived optimal thresholds for comparison, which in all cases was well within the range of optimal performance. (B) Hit ratios for different load parameters, depicted for various levels of *p* (see legend). All models show optimal performance with a hit ratio of 1 for low values of *α*, but undergo a fast transition to a non-retrieval network at different, higher load values. (C) Histograms of Hamming distances for *p* = 5% (left panel) and *p* = 50% (right panel) and for five different load parameters around the transition from a fully functioning to a non-retrieval network. (D) 90% percentile of Hamming distance for different load parameters and for *p* = 5% (left) and *p* = 50% (right). The red dashed lines show the distances and load parameters of the points with maximum curvature.

### 3.2 Network capacity

To determine the capacity of the networks we examined the hit ratio as a function of the load parameter for different levels of pattern bias. We found that as the pattern bias decreased, the transition point from a hit ratio of 1 to 0 occurred for higher values of the load parameter, meaning that the network’s capacity increased (Figure 3B). The transition from high to low hit ratios also became slower for the sparser patterns (see e.g. *p* = 2%). Only the 1% sparsely coded networks could consistently store more patterns than the number of nodes, i.e. *α >* 1 still yielded hit ratios of 1. The capacity of the network for unbiased patterns was between 0.13 and 0.15, in agreement with the results from the replica theory (Amit et al., 1985b). The higher load capacity for sparse patterns, as confirmed by the numerical simulations, points to the question whether this higher capacity also translates to a higher robustness of biased patterns more generally.

Before looking at robustness, we first studied the behavior of networks at the edge of their breakdown in performance in more detail. To do so, we examined the distributions of Hamming distances between the desired memory patterns and the settled stable states for the cases of *p* = 5% and *p* = 50% (Figure 3C). For load parameters considerably below the critical value *α*_*c*_, all simulations produced zero or negligible Hamming distances, indicating successful retrievals. However, when surpassing the network’s capacity, the stored patterns were not anymore the point attractors of the network and large distances between the memory patterns and the settled states emerged. The shifts between these two modes of the histograms were very rapid (compare e.g. *α* = 0.11 and *α* = 0.17 in Figure 3C).

To capture the behavior of the network as the loading exceeds the capacity, we took the 90-th percentile of Hamming distances for a varying load *α* for *p* = 5% and *p* = 50% as examples (Figure 3D). For each we found an abrupt jump in the Hamming distances, confirming a sudden breakdown in performance. The Hamming distance from which a quick departure took place was considerably higher for unbiased patterns. This indicates some corruption of the pattern retrieval occurring already before the overall performance breakdown, potentially related to a difference in robustness between unbiased and bias patterns.

### 3.3 Measuring performance and robustness

To determine how different types of noise affected the load capacity and hit ratio depending on the pattern bias, we used a robustness measure that incorporated a sparseness-dependent tolerance based on mutual information (see Methods). The robustness measure thereby provided an aggregated measure of the network’s behavior, which we evaluated for each noise type, as the normalized AUC of the hit ratios (Figure 4).

**Figure 4:**
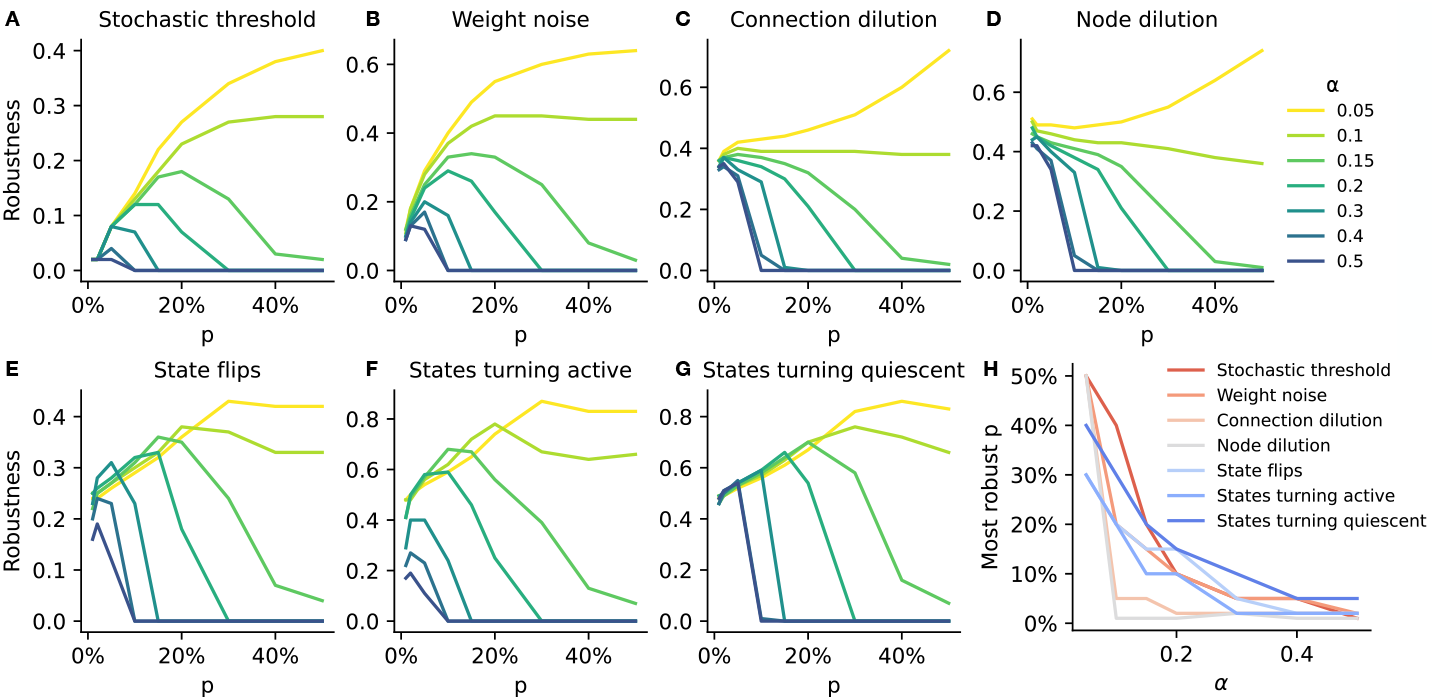
Robustness of the Hopfield networks against various kinds of disruptions. (A-G) Each panel shows the robustness measure for different pattern bias and load parameters (colored lines) against the indicated noise type. (H) Aggregated results indicating the most robust pattern bias for each pattern load and noise type.

In the simulations with a stochastic threshold, equivalent to noise from the temperature parameter, unbiased patterns (*p* = 50%) had the highest robustness for settings with a low pattern load (*α* = 0.05; Figure 4A). In contrast, for high pattern loads (*α* ≥ 0.4), networks with both biased and unbiased patterns exhibited very low robustness. However, at intermediate levels of pattern load, networks using intermediate levels of pattern bias were most robust. This indicates that the pattern load that a networks receives determines which pattern representation is most suitable for retrieval of memory patterns in light of noise.

The simulations using weight noise yielded very similar results (Figure 4B), with the only difference being a slight shift in the robustness values, while the relation between pattern load and robustness of the pattern bias remained the same. Therefore, both types of noise, unbiased patterns were only robust for low pattern loads, while biased patterns were more robust with increasing pattern loads.

The noise types involving connection and node dilution yielded overall qualitatively similar results, even though sparser networks were more robust here in general, irrespective of the pattern load (Figure 4C&D). Therefore, there was a more abrupt shift, so that a slight increase in pattern load lead to the sparse patterns being most robust.

For the three noise types that involved changes in the patterns presented during retrieval (state flips, states turning active, and state turning quiescent), sparse patterns again exhibited a robustness advantage for higher loads (Figure 4E-G). With a gradual increase in pattern load, there was a gradual increase in the pattern bias that was most robust.

To summarize the relation between the pattern load of a network with the pattern bias that performs most robust against different types of noise, we determined for each pattern load the pattern bias *p* that was associated with the highest robustness. For example, for the stochastic threshold, *α* = 0.15 had the peak robustness for a mean activity *p* = 20%. The aggregated results illustrated that with increasing pattern loads, an increase in patterns bias ensures the most robust performance (Figure 4H). For connection and node dilution, there is an abrupt change in more robust performance of biased patterns. For the other noise types the change is more gradual, so that intermediate levels of pattern bias perform most robust for intermediate pattern loads.

We conclude that sparse patterns can be more robust against different types of noise compared to unbiased patterns in Hopfield networks. Importantly, for most robust performance, the pattern bias should be chosen according to the expected pattern load.

## 4 Discussion

We studied the robustness of pattern retrieval in Hopfield networks for seven different types of biologically-motivated noise. To do so, we first analytically determined optimal thresholds and tolerance levels for different levels of pattern bias. Then we studied the retrieval performance as a function of the pattern load and the pattern bias to see what happens in the regime where the retrieval performance breaks down. Finally, we applied a robustness measure, which aggregates the hit ratio over a range of noise strengths, and found that different pattern loads required different levels of pattern bias for most robust performance. Overall, the higher the pattern load, the more biased the most robust pattern representation had to be.

To measure the pattern retrieval performance, we defined the hit ratio based on a corruption tolerance. This is more appropriate than using e.g. the mean Hamming distance because the Hamming distance of a failed retrieval from the stored pattern depends on the pattern bias. For example, in the case of connection dilution, after deleting many connections, an entirely quiescent pattern becomes the network’s stable state. This non-retrieval state has on average a Hamming distance *p* to the stored patterns. In contrast, the expected distance between two random patterns in the network is 2*p*(1−*p*), meaning that the retrieval of a mistaken memory results in lower Hamming distances for biased patterns. To mitigate this issue for comparing performance of different levels of pattern bias, we derived the tolerance levels based on mutual information. This approach was further supported by our finding that the Hamming distances before the abrupt degradation in the retrieval performance (Figure 3D) agreed with our differentiated choice of tolerance values.

While we found that different types of noise exhibit a similar relation between the pattern load and the most robust pattern bias (Figure 4H), the underlying mechanisms and dynamics are likely to be more complex and heterogeneous across the different noise types. For the stochastic threshold noise type, sparse patterns had very low robustness even for low pattern loads (Figure 4A). Intuitively, this is plausible because for sparse patterns a stochastic threshold means that the activation of the few active nodes is at risk of being changed and therefore the pattern might not be retrieved. More formally, less biased (i.e. more distributed) patterns have a higher information load with the total entropy of the stored patterns being proportional to the Shannon function *h*(*p*) = −*p* log_2_(*p*) − (1 − *p*) log_2_(1 − *p*) (Frolov et al., 1997), which is low for sparse patterns and maximal for the unbiased patterns. For Hopfield-like memory networks in the brain this suggests that activity fluctuations of single neurons that are part of sparse patterns are detrimental for accurate pattern retrieval. Possible ways to reduce the risk for this detrimental effect in the brain include a low, stable baseline activity in single neurons of these networks. Some evidence supporting this possibility comes from the observation that neurons in the medial temporal lobe with less selective responses to sensory stimuli have higher baseline firing rates than neurons with more selective responses (Mormann et al., 2008). Furthermore, medium-spiny neurons in the striatum, a region associated with sparse coding of actions, exhibit very low baseline activity (Berke et al., 2004; Markowitz et al., 2018).

For our simulations with weight noise, sparser representations slightly improved in their robustness compared to the stochastic threshold noise type (Figure 4B). Thus, the mean activity level of the most robust networks against weight noise was lower than that in the stochastic model (Figure 4H). However, as the range of the noise values was chosen to cover the network’s behavior from a successful retrieval to a breakdown, the robustness values for different noise types are not directly comparable. Instead they indicate within a given noise type the relative performance of different pattern representations and pattern loads. For weight noise, perturbations in the connection strengths cause the weight matrix to lose its symmetry and thereby a sufficient condition for the existence of the energy function, even though such disturbed networks can still operate as attractor systems (Rolls & Deco, 2010). After the introduction of asymmetry in the weight matrix, the mean field theory can be applied for the case of *M* ≪ *N*, and it was shown that asymmetry is then analogous to increasing the temperature in the network (Hertz et al., 1986) and can make spurious states unstable, potentially improving the network’s performance (Hertz et al., 1991). Possibly, this connection underlies the qualitatively similar retrieval for the two noise types and accounts for their similar robustness profiles (Figure 4A&B). One limitation is that we only considered static weight noise here, so that a single post-learning perturbation took place in the connections. In general, the brain is exposed to more dynamic noise, e.g. for weight noise through synaptic strengths (Ribrault et al., 2011), which affects the network during learning and retrieval with varying perturbations.

Similar to weight noise, also connection dilution introduces an asymmetry in the weight matrix. Interestingly, this had a different effect though on the robustness in relation to pattern bias (Figure 4C). Overall, biased patterns had high robustness with peak performance for sparse patterns and *α >* 0.1. It seems that here, next to the information load (see above), a second, opposing dynamic exists which favors sparsely-coded patterns and makes them less vulnerable to spurious minima, increasing their load capacity. This dynamic could be related to the reduction of spurious states in asymmetric networks (Hertz et al., 1991). However, in our simulations the asymmetry was introduced by a random, post-learning dilution of the connections, which is different from a network without all-to-all connectivity, or a network with a learning rule for asymmetric connection weights. In the brain, connection dilution (or synaptic loss) strong correlates with cognitive decline (Scheff et al., 2006). Horn et al. (1993) used a similar Hopfield network to investigate the deterioration of memory retrieval due to synaptic loss in Alzheimer’s disease. The random deletion of connections by the probability *d* corresponds to multiplying the connection strengths *W*_*ij*_ by the Bernoulli variables *d*_*ij*_, which are 1 with the probability (1 − *d*), and 0 otherwise. Considering the signal-to-noise ratio, this means that the two signal values (the mean values of the nodes’ local fields for the 0 and 1 states), before introducing an activation threshold, decrease proportional to (1 − *d*). Thereby, the threshold is no longer optimal, i.e. keeping the local fields in a situation similar to Figure 2A instead of Figure 2B. Thus, the signal-to-noise ratio reduces and performances declines, providing a potential explanation for some of the cognitive deficits in Alzheimer’s disease.

The results from simulations involving node dilution showed a strong similarity to connection dilution (Figure 4C&D). For load parameters smaller than capacity, the robustness levels were far from zero in both cases, indicating that a memory breakdown happened only after a high number of deletions. Based on the higher capacity of sparse pattern representations in Hopfield networks, one could expect that they are in general also more robust. However, our results demonstrate that this is not always the case. This can be seen e.g. in the effect of node dilution on the signal-to-noise ratio. The total number of nodes is reduced by the node loss, but the number of patterns remains constant. This change is different from simply an increase in the load parameter, since the disruption takes place after the learning phase. Using the optimal uniform threshold *θ*_*opt*_ from Eq. 8, the signal-to-noise ratio for one bit of a stored pattern is given by Eq. 9. For a deletion ratio *d*, the signal and cross-talk noise terms in Appendix B must be adjusted such that the number of summed terms and thus both signals (before threshold) and cross-talk noise are reduced by the factors (1 − *d*) and 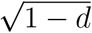, respectively. Therefore, the signal-to-noise ratio for the model with threshold *θ*_*opt*_ and deletion ratio *d* is given by:

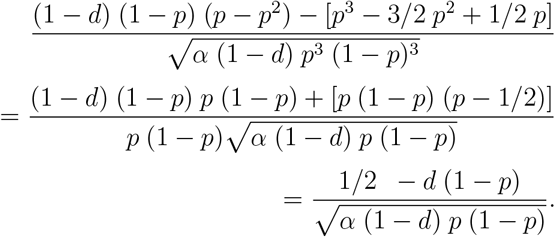

Therefore, the deterioration in the signal-to-noise ratio after deletion of nodes is stronger for sparser patterns, especially for higher values of *d*, which indicates that the higher information load in more distributed patterns can contribute to robustness. This is in line with the view that ‘local codes’, i.e. very sparse patterns here, are sensitive to error and noise, and the redundancy in more distributed representations improves fault tolerance (Spanne & Jörntell, 2015). Nevertheless, in our simulations, for increased pattern loads, the robustness advantage of sparse patterns relative to more distributed patterns was more pronounced for connection and node loss in comparison to stochastic thresholds and weight noise. This indicates that sparse patterns in Hopfield-like networks in the brain might be useful to reduce their vulnerability to structural loss in brain regions such as hippocampus and thereby postpone the onset of cognitive decline e.g. in Alzheimer’s disease.

The other types of noise we examined manifested as distorted versions of the stored patterns presented during the retrieval phase, i.e. states were made active, quiescent, or flipped their values (Figure 4E-G). The results between these types of noise were qualitatively similar. However, the advantage of sparse patterns was strongest for the case of states turning quiescent (Figure 4G), since there was a smaller probability of the few active units to be affected by that change. The high robustness values for states turning active or quiescent point to successful retrieval even for a large number of activated or silenced units (Figure 4F&G). In addition the robustness profiles exhibited pronounced peaks for each pattern load, providing a specific level of pattern bias that performed most robust. This points to another potential advantage of sparse pattern representations in the brain, so that additional activation of neurons that do not belong to the stored pattern, does not strongly impair the pattern retrieval. For example, a population pattern in the hippocampus represented by sparse place field activity would be robust against conflicting activation of random place cells. This type of conflicting activation (or deactivation) might also mimic the application of optogenetics in neuroscience in which neuronal activity of a targeted part of a neural circuit can either be activated or silenced by using light with a millisecond temporal resolution (Boyden et al., 2005). In contrast, state flips as a type of noise when presenting a pattern for retrieval entails knowing the actual memory pattern and then distorting it, which may not have an obvious biological correspondence.

Our results on the relation between pattern bias and robustness are relevant for the ongoing discussion about coding schemes and pattern representation in the brain. Different brain regions seem to employ different coding schemes in the range from sparse to distributed representations (Quiroga & Kreiman, 2010). This might reflect different functional specializations as coding schemes relate to the ability to generalize (Spanne & Jörntell, 2015). For example, neural coding in the hippocampus, with the very localized activity of place cells, is considered to be sparse, while cortical regions are often associated with more distributed representations (Quiroga & Kreiman, 2010). Based on our results the question of which coding scheme is optimal for a given brain region, should also consider the expected pattern load. Intuitively, a given memory network in the brain should have a capacity that is as high as possible. However, there are many constraints on memory systems such those imposed by sparseness on memory lifetimes (Leibold & Kempter, 2007) and the ability to store sequences (Leibold & Kempter, 2006), which are arguably just as important as capacity. Furthermore, Hopfield-like networks in the brain might be used to store patterns more generally, i.e. not necessarily only patterns related to semantic or episodic memories, but also non-declarative memories. On an evolutionary time scale, these networks might have been optimized to a specific level of sparse or distributed representations that is the result of a trade-off between capacity, robustness, and the expected pattern load. For example, a brain network that is typically only used for a low pattern load, might be able to afford a rather distributed pattern representation, providing high robustness against various types of noise (Figure 4H). In contrast, networks that are used to store a high number of patterns might have evolved to use a more sparse pattern representation, again providing high robustness against various types of noise. This view is in line with the proposal that sparse representations are used by memory systems to achieve a high capacity, while distributed representations are used by sensory systems (Rolls & Treves, 2011). However, our results emphasize that not only the extremes of the coding schemes, i.e. sparse or fully-distributed, should be considered, as for certain intermediate pattern loads, also intermediate levels of pattern bias might be advantageous.

It has been pointed out several times that the current use of the terminology related to pattern representations and neural coding is inconsistent across different subfields (Quiroga & Kreiman, 2010; Spanne & Jörntell, 2015; Beyeler et al., 2019). We used the term *unbiased* here to refer to patterns with *p* = 0.5, which would correspond to a fully distributed (Rolls & Treves, 2011) or dense (Spanne & Jörntell, 2015) representation. Biased patterns (here *p <* 0.5) are also referred to as sparse distributed representations (Rolls & Treves, 2011), while we used the term *sparse* to indicate that *p <<* 0.5 and the term *sparseness* to refer to the degree to which the representation deviates from *p* = 0.5.

While Hopfield networks are an established model for memory networks in the brain, they make strong abstractions of the underlying biological networks and their dynamics. For our simulations these abstractions are important to consider for the effects of noise on different pattern representations. Firstly, the patterns we examined here were uncorrelated, i.e. for each pattern the active and inactive units were randomly selected. In contrast, neural activation patterns are often correlated, e.g. due to the spatial activity patterns of hippocampal place cells, which is problematic for Hopfield-like storage of memory patterns e.g. in CA3 (Papp et al., 2007; Neher et al., 2015). Secondly, the symmetric all-to-all connectivity in Hopfield networks does not match connectivity rules of biological neural networks or cell-type specific connectivity. For these reasons hippocampal models using Hopfield networks are often modified to better account for neurophysiological characteristics (Rolls, 2013). While our results on the robustness consider here only a general type of Hopfield network, it remains to be shown how neurophysiological characteristics of specific brain regions and networks affect robustness. Finally, although we used a relatively large network (*N* = 5000), the properties of sparse and distributed codes, including robustness, might be affected by the network size (Spanne & Jörntell, 2015). Despite these limitations our results suggest that the performance of Hopfield networks should not be judged entirely based on capacity considerations, but also based on the robustness against different types of noise, which are relevant for biological neural networks. Our findings indicate that the coding schemes employed by different brain regions are adapted so that their vulnerability to disruptions is mitigated.

## Appendix

### A) Signal-to-noise ratio analysis in the modified Hopfield network

Assumed is a uniform activation threshold *θ*. Starting from the update rule for the state of node *i*, we have

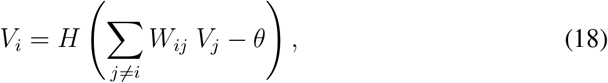

where *H* is the Heaviside step function.

Using the Eq. (5), we expand the weights in Eq. (18):

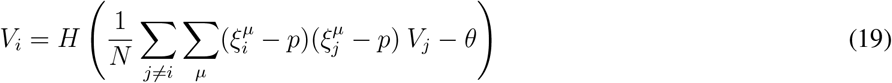

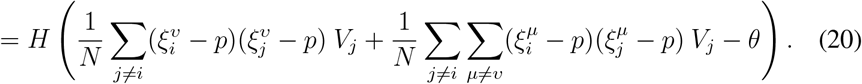

When the network is in the stored pattern *υ*, it means that 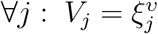, thus

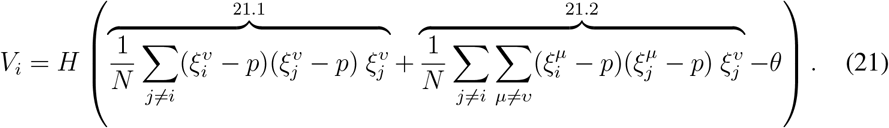

Since 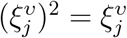 the expression 21.1 is equal to

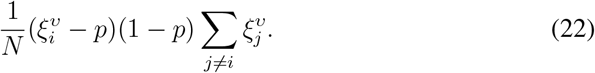

Due to the independence of the memory bits, we have

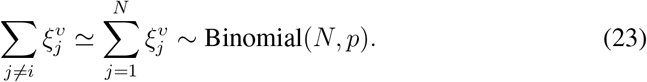

Hence,

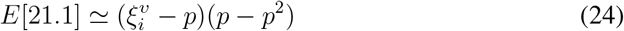

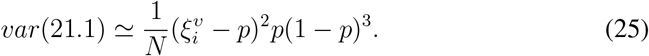

On the other hand, the expression 21.2 consists of (*M* − 1)(*N* − 1) terms, each with the following possible outcomes and probabilities:

Hence,

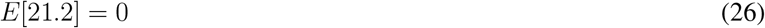

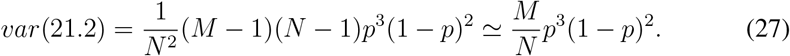

We express results in probabilistic terms, since there is a large number of elements drawn from the same distribution. In reality, the results are deterministic and based on the realizations of the Bernoulli variables.

If the expressions 21.1 and 21.2 are independent and by ignoring the smaller variance in 21.1, we can approximate the total terms inside Eq. (21) as the distributions summarized in Table 2, based on the *ξ*-pattern state value 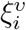.The optimal threshold *θ*_*opt*_ is equalizes the absolute values of the two means, as shown in the Figure 2B.

**Table 1:** Probability table of the elements in the expression 21.2.

| Outcome | Probability |
| --- | --- |
| 0 | $1 - p$ |
| $p^2$ | $(1 - p)^2 p$ |
| $p^2 - p$ | $2p^2(1 - p)$ |
| $(1 - p)^2$ | $p^3$ |

**Table 2:** Signals and cross-talk noises for different states of a memory pattern.

| State | Mean | Variance |
| --- | --- | --- |
| $\xi_i^v = 0$ | $-p(p - p^2) - \theta$ | $\alpha p^3(1 - p)^2$ |
| $\xi_i^v = 1$ | $(1 - p)(p - p^2) - \theta$ | $\alpha p^3(1 - p)^2$ |

**Table 3:** Probability table of the elements in the expression 28.2.

| Outcome | Probability |
| --- | --- |
| $-p^3$ | $(1 - p)^3$ |
| $p^2(1 - p)$ | $3(1 - p)^2 p$ |
| $-p(1 - p)^2$ | $3(1 - p)p^2$ |
| $(1 - p)^3$ | $p^3$ |

### B. The reduced cross-talk noise by adding correlated random local fields

By a transformation in form of *V*_*j*_ → *V*_*j*_ − *p*, Eq. (21) changes to

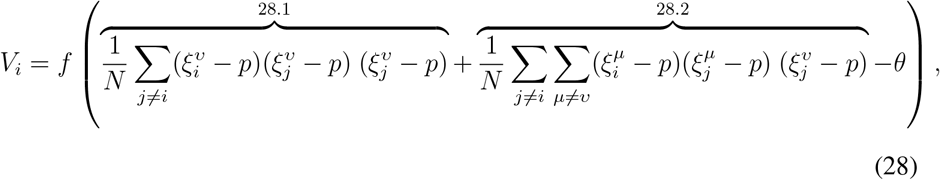

where the expression 28.1 is equal to:

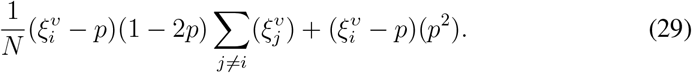

Using Eq. (23), we have

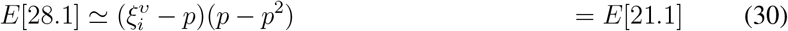

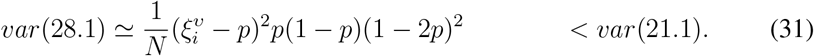

The mean and variance expressions above show that the signal is unchanged.

Regarding the expression 28.2, there are again (*M* − 1)(*N* − 1) terms, each with the following possible outcomes and probabilities:

Hence,

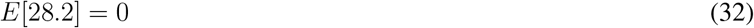

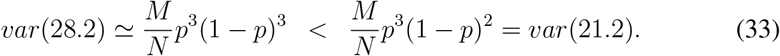

The decreased variance of 28.2 in comparison to 21.2 means that the transformation *V*_*j*_ → *V*_*j*_ − *p* or equivalently adding the random thresholds *p*Σ _*j ≠i*_ *W*_*ij*_ decreases the noise term.

## Acknowledgments

We thank Heinz-Jürgen Schmidt for interesting discussions and helpful comments on a previous version of this manuscript.

## Code availability

The code for the simulations done in this manuscript and their visualization is available here: https://gitlab.ruhr-uni-bochum.de/smidtrc8/robustness-in-hopfield-networks.

